# SwiftNJ: Fast Exact Neighbour Joining via Correctness-Gated Coding Agents

**DOI:** 10.64898/2026.05.28.728410

**Authors:** Joe Christensen

## Abstract

The capability profile of frontier coding agents in 2026 varies sharply across technical domains, motivating domain-specific empirical study of where, and under what oversight conditions, such systems can contribute to specialised technical work. This paper presents one such study in computational phylogenetics. Neighbour joining (NJ) is a widely used distance-based method for inferring evolutionary trees in microbial epidemiology, comparative genomics, and large-scale sequence clustering. Its constant-factor runtime is set by hand-tuned native implementations; RapidNJ is a widely-cited representative of that class and serves here as the comparison baseline. We ask whether a current-generation coding agent, operating under a correctness-gated optimisation harness with deterministic correctness gates calibrated against a QuickTree reference, can advance that constant factor on a fixed benchmark. The resulting implementation, SwiftNJ, achieves a geometric-mean runtime ratio of 0.565 against a locally-rebuilt RapidNJ-native binary across a 59-matrix corpus, sub-parity on 58 of 59 matrices. On 400 shuffled inputs drawn from 16 small matrices (*n* ≤ 2000), SwiftNJ matched the QuickTree reference at Robinson–Foulds distance zero. In this domain, a correctness-gated coding agent meaningfully improved on a strong native baseline, suggesting that harness-guided optimisation holds promise for performance-critical bioinformatics tools; further work is needed to establish how broadly the approach generalises.

## 1 Introduction

The capabilities of frontier AI coding agents are advancing on the timescale of months, and the resulting capability landscape is uneven. Field-experimental evidence from Dell’Acqua et al. [1] characterises this landscape as a *jagged technological frontier*, with large measured gains on tasks inside the frontier and measurable losses on tasks outside it, and with the boundary irregular and difficult to predict from intuition. Empirical exploration of where these systems may hold promise is therefore an important task in determining whether, and where, they can be useful in scientific work.

Recent developments in mathematics [2, 3, 12, 11] and software security [4, 5, 6] suggest that this frontier is moving quickly. Comparable demonstrations on scientific software in the life sciences are by comparison sparse, leaving open the empirical question of whether agentic coding workflows can advance bioinformatics targets where native implementations already define the state of the art.

This paper presents one such empirical probe in computational phylogenetics. Exact neighbour joining [7] is a foundational distance-based method in the life sciences, with applications in microbial taxonomy, viral phylogenetics, and large-scale sequence clustering. Its constant-factor state of the art is held by hand-optimised native implementations, of which RapidNJ [8] is representative: a C++ implementation built on per-row lower bounds on the *Q* criterion that skip work without affecting the selected join.

Exact NJ is a good test case for this question: it has a short kernel, a precise correctness oracle, and cheap repeated measurement, all of which let a harness make iteration-level keep-or-revert decisions cleanly. We therefore use it to probe whether a harness deploying current-generation frontier coding agents, Opus 4.7 (xhigh) in Claude Code and GPT-5.4 (xhigh) in Codex (where xhigh denotes the reasoning-effort setting), can produce a useful artefact. RapidNJ-native serves as the comparison baseline. A single Opus 4.7 (xhigh) instance acts as the running agent with write access only to the Rust source. We supervised the loop directly, watching for mode collapse and adjusting the corpus, gates, and harness as the search progressed; the two further agents in Claude Code and Codex served as collaborators in this oversight rather than as autonomous maintainers. Each iteration is accepted or reverted by a hard correctness gate (Robinson–Foulds distance zero against a reference QuickTree implementation at *n* ≤ 2000, branch-length tolerance at larger *n*) and a per-regime geometric-mean runtime ratio score against RapidNJ-native.

The resulting implementation, SwiftNJ, is faster than a locally-rebuilt RapidNJ-native binary on 58 of 59 benchmark matrices and on all five matrices in a separate large real-derived transfer check.

## 2 Background

Neighbour joining [7] is one of the most widely used distance-based methods for phylogenetic inference. The canonical algorithm is *O*(*n*^3^). RapidNJ [8] reduced the constant factor by introducing per-row lower bounds on the *Q* criterion that skip work without affecting the selected join, and remains among the fastest published exact NJ tools. DecentTree [9] packages NJ and its variants in a unified, performance-tuned C++ implementation. QuickTree [10] is a widely-cited canonical NJ implementation used here as the reference for the small-matrix correctness gate at *n* ≤ 2000, where its Robinson–Foulds distance [13] is taken against SwiftNJ’s output.

The methodology used in this work is parallel in spirit to Karpathy’s autoresearch [16], which iterates on a language-model training script under a similar keep-or-discard regime. The principal differences here are the target domain and the form of the correctness oracle: rather than a learned evaluator or an LLM-as-judge, the gate is a deterministic comparison against a reference tree.

## 3 Method

### 3.1 Roles

The harness comprises a roughly 400-line Python orchestrator and a fixed benchmark corpus, with three roles distinguished:

- **Operator** (me). Defined the task, the corpus, the correctness gates, and the scoring rule. Supervised the loop in real time: watching for mode collapse, evaluating corpus coverage and gate calibration, deciding when to revert to an earlier accepted point, and adjusting the harness as the search progressed.
- **Running agent**. A single Opus 4.7 (xhigh) instance with write access only to the SwiftNJ Rust source. Its input is the current source, the recent iteration log, and the current score. It never reads or writes the harness, the correctness gates, or the scoring rule.
- **Collaborator agents**. An Opus 4.7 (xhigh) instance in Claude Code and a GPT-5.4 (xhigh) instance in Codex, used by the operator for writing and modifying harness Python, evaluating proposed gate changes, and providing a second opinion on judgement calls. These agents did not autonomously maintain the loop; they implemented the operator’s decisions and served as sounding boards on ambiguous iterations.

### 3.2 Iteration loop

Each iteration proceeds as follows. (1) The running agent receives the current Rust source and the recent iteration log. (2) It proposes a patch. (3) The harness applies the patch and runs `cargo build --release`. (4) For each matrix in the visible corpus, the harness parses the Newick output and verifies that the taxon set matches; at *n ≤* 2000, that the Robinson–Foulds (RF) distance against a cached QuickTree reference is zero; at *n >* 2000, that the total branch length matches a stored reference within tolerance. (5) Runtime is measured as the median of repeated single-threaded runs (`RAYON_NUM_THREADS=1`). (6) The score is the geometric mean across regimes of the per-matrix runtime ratio against the RapidNJ-native baseline.

A patch is kept if and only if it passes all correctness gates and improves the score. Otherwise it is reverted. There is no partial credit for an interesting-looking patch that regresses the score, and no acceptance of any patch that fails a correctness gate, regardless of speedup.

### 3.3 Accepted algorithm

The candidate accepted at iteration 51 combines five ingredients, each introduced by a specific kept iteration:

- **Fixed-stride matrix layout (iter 2)**. Distances are stored in a contiguous `f32` buffer with a fixed power-of-two stride. SIMD loads are aligned and the next-row pointer is a constant offset.
- **AVX2 running-min** *Q* **scan (iter 2)**. Wide *f* 32 *×* 8 inner loop with a running-minimum register, replacing the scalar baseline.
- **Row-level** *Q* **pruning (iter 5)**. A per-row lower bound on *Q*; rows that cannot improve on the current best *Q* are skipped. This is the RapidNJ row-bound idea applied to the new layout and represents the first sub-parity iteration.
- *f* 16 **chunk-min pruning (iter 27)**. Per-chunk minima stored in half precision, rounded down on store so the stored value remains a conservative lower bound, halving the bound-check bandwidth.
- *O*(1) **dead-taxon skip (iter 43)**. Joined taxa are excised from the active set in *O*(1) via an active-index permutation rather than by rebuilding the matrix.

Each lower bound is provably no greater than the true *Q* value of the rows or chunks it covers. A skip is taken only when the bound rules out improvement on the current best *Q*; a loose bound therefore costs performance, not correctness. The exactness claim is under the benchmark input contract: finite, nonnegative, symmetric PHYLIP distance matrices with zero diagonal and no NaN or Inf values. Conservative lower-bound rounding can only cause extra scanning, never an incorrect skip.

### 3.4 Corpus and machine

The corpus has 59 distance matrices in five regimes: 14 matrices from a course assignment at the Bioinformatics Research Centre (89 to 1849 taxa, “Course”), synthetic matrices from five deterministic generator regimes (uniform-random, additive-Yule, clustered-blocks, redundant-rows, and concentrated-distances) at *n ∈{*1000, 2000, 5000, 10000*}*, and real Pfam-derived matrices [14]. Of the 59, 37 were visible during the loop and entered the score; 22 were held out and never seen by the running agent.

All comparisons are matrix-to-tree only; alignment-to-distance work is excluded for both tools. All runs are single-threaded. SwiftNJ and RapidNJ-native are both built locally with `-march=native`; RapidNJ-native is built from upstream sources without modification. The machine is an AMD Ryzen 5 5600H (6 cores, 12 threads, AVX2/F16C/FMA, no AVX-512) running Windows 11 with WSL2 Ubuntu 24.04, Rust 1.95, GCC 13.3. Per-matrix runtimes are medians over ten repeated runs.

## 4 Results

### 4.1 Per-regime runtime ratios

Figure 1 summarises the headline result. SwiftNJ is below the RapidNJ-native parity line in every regime. Per-regime geometric-mean ratios appear in Table 1. DecentTree’s NJ-R-V and NJ-V modes are both slower than RapidNJ-native on this benchmark and machine; they are included for context.

**Table 1:** Per-regime geometric-mean runtime ratio of SwiftNJ against RapidNJ-native and the implied speedup.

| Regime | Ratio | Speedup |
| --- | --- | --- |
| Course (89–1849) | 0.644 | 1.55× |
| Synthetic 1k–2k | 0.488 | 2.05× |
| Synthetic 5k | 0.547 | 1.83× |
| Synthetic 10k | 0.713 | 1.40× |
| Real Pfam | 0.465 | 2.15× |
| All 59 matrices | 0.565 | 1.77× |

**Figure 1:**
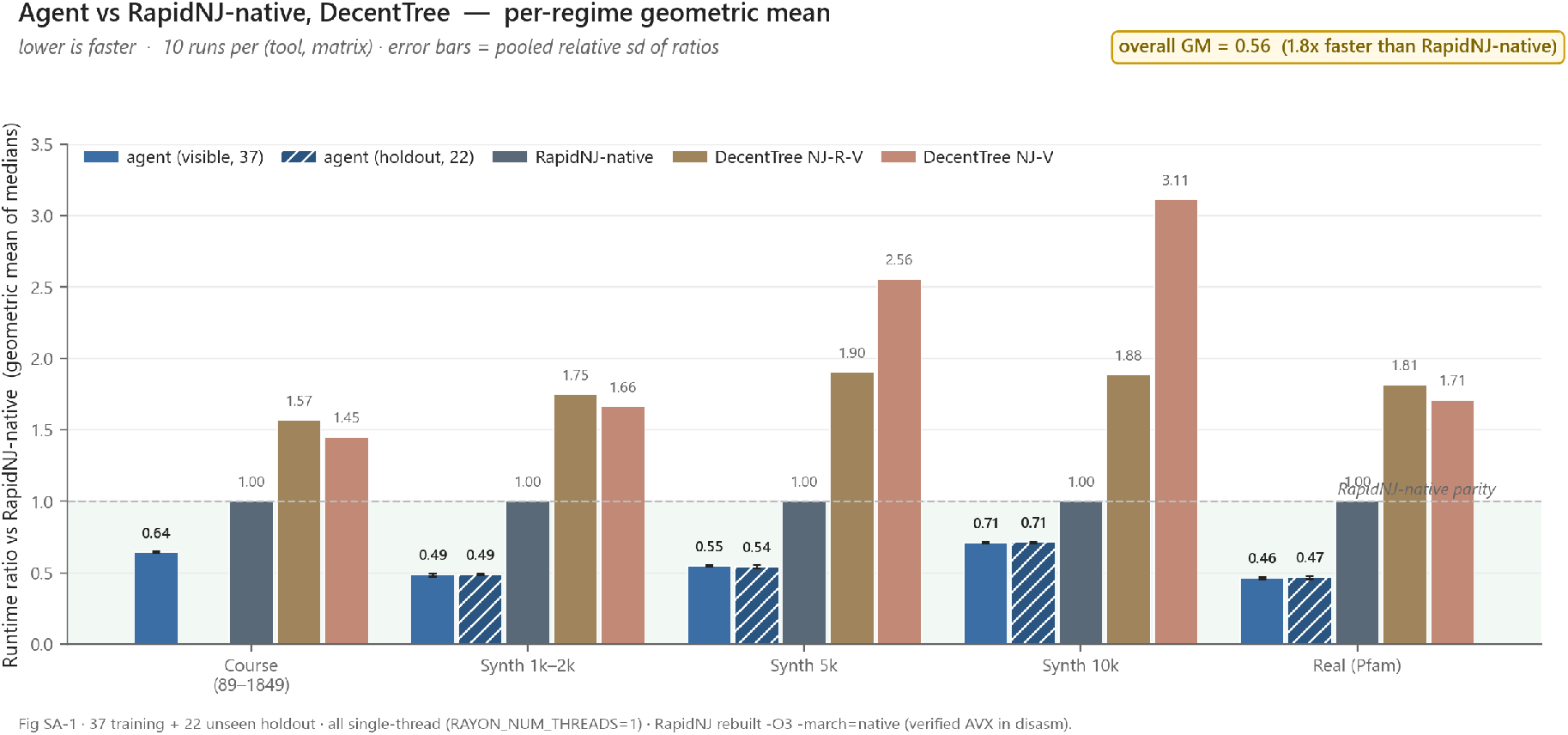
Per-regime geometric-mean runtime ratio of SwiftNJ against RapidNJ-native. Lower is faster. DecentTree (NJ-R-V and NJ-V) is shown for context. Across all 59 matrices the geometric-mean ratio is 0.565, sub-parity on 58 of 59 matrices; the worst-case ratio is 1.005 on a single small matrix.

### 4.2 Visible versus holdout

Figure 2 compares visible and holdout per-regime geometric means. The two overlap to within 0.01 in every regime, indicating that the speedup does not collapse on matrices the running agent had not seen during the loop. The Course regime is visible-only and is therefore omitted from the holdout comparison.

**Figure 2:**
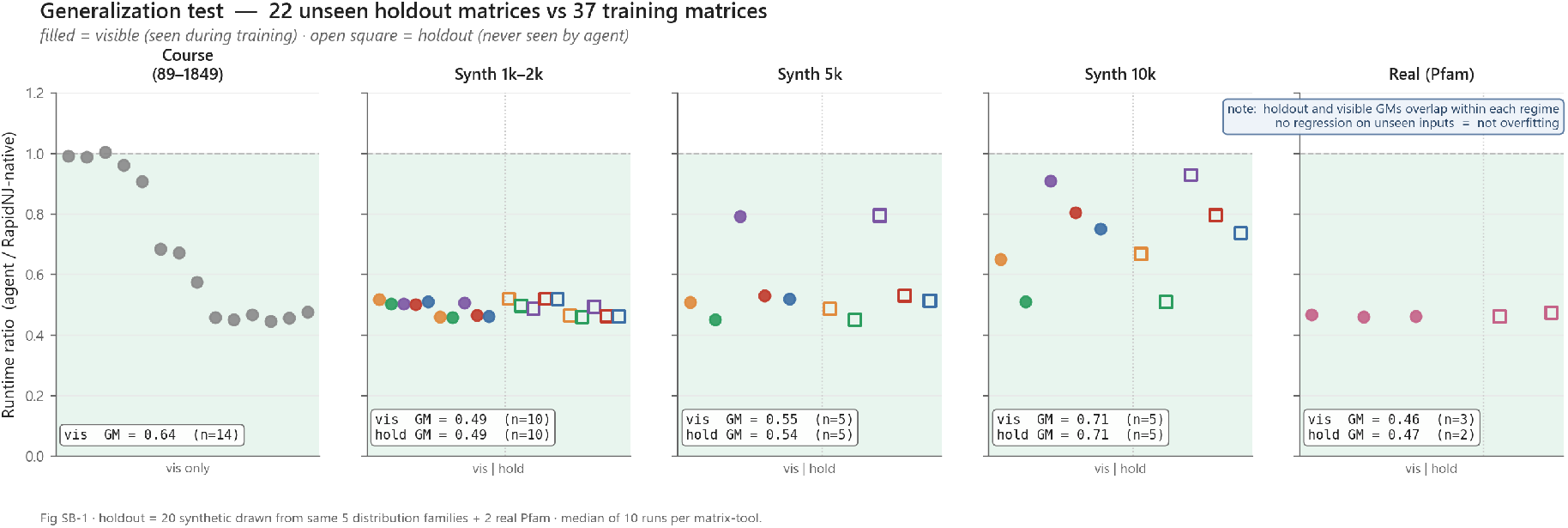
Visible (closed markers) versus holdout (open markers) per-regime geometric-mean runtime ratios. Closed and open markers overlap to within 0.01 in every regime.

### 4.3 Large real-derived matrices

A separate transfer check used five real or real-derived biological distance matrices spanning 4,986 to 10,000 taxa: prokaryotic ribosomal RNA alignments from the Comparative RNA Web Site [17] (CRW 16S and 23S) and clustered viral hemagglutinin alignments (H1N1pdm, H5N1) derived from public GenBank/INSDC records via GenSpectrum/Loculus [15]. None of the five matrices entered the optimisation loop. SwiftNJ was nominally faster on all five, with four clear wins and one near-parity case (H5N1, 1.05 *×*), for a geometric-mean speedup of 1.43 *×* (Table 2). These *n >* 2000 cases were not RF-adjudicated by QuickTree; checks were limited to parser and taxon-set consistency and to branch-length and timing comparisons.

**Table 2:** Large real-derived matrix transfer check. Median over 10 repeats per tool, matrix-to-tree only, single-threaded, same machine.

| Case | Taxa | SwiftNJ (s) | RapidNJ-native (s) | Speedup |
| --- | --- | --- | --- | --- |
| CRW 16S bacteria prefix | 10,000 | 18.40 | 34.94 | 1.90× |
| CRW 16S three-domain random | 6,000 | 5.42 | 8.21 | 1.52× |
| CRW 23S bacteria | 4,986 | 3.26 | 4.72 | 1.45× |
| H1N1pdm HA | 8,000 | 11.27 | 15.51 | 1.38× |
| H5N1 HA clade 2.3.4.4b | 8,000 | 9.85 | 10.36 | 1.05× |
| Geometric mean |  |  |  | 1.43× |

### 4.4 Shuffle invariance

To verify that the speedup is not a tie-breaking artefact of a particular row ordering, 16 matrices were each run with 25 random row permutations (400 shuffled inputs in total) and SwiftNJ’s output was compared against the QuickTree reference by RF distance. All 400 matched (RF distance zero). Direct RF equality between SwiftNJ and RapidNJ is not claimed: RapidNJ and QuickTree return different but equally valid NJ trees on some matrices because of tie-breaking convention, and at *n ≤* 2000 SwiftNJ and RapidNJ disagree in the same places that RapidNJ and QuickTree disagree.

### 4.5 Search trajectory

Figure 3 shows the trajectory of accepted scores across the 51 iterations. Iteration 1 was a faithful Saitou–Nei port at approximately 3.0*×* RapidNJ-native runtime. The agent then introduced, in order, the AVX2 running-min *Q* scan (iter 2), row-level *Q* pruning (iter 5, first sub-parity), *f* 16 chunk-min pruning (iter 27), and the *O*(1) dead-taxon skip (iter 43). Of 51 iterations, 25 were kept and 26 rejected (keep rate approximately 49%). The final harness score was 0.61; a clean re-measurement, with measurement drift controlled, gave a corrected score of 0.565.

**Figure 3:**
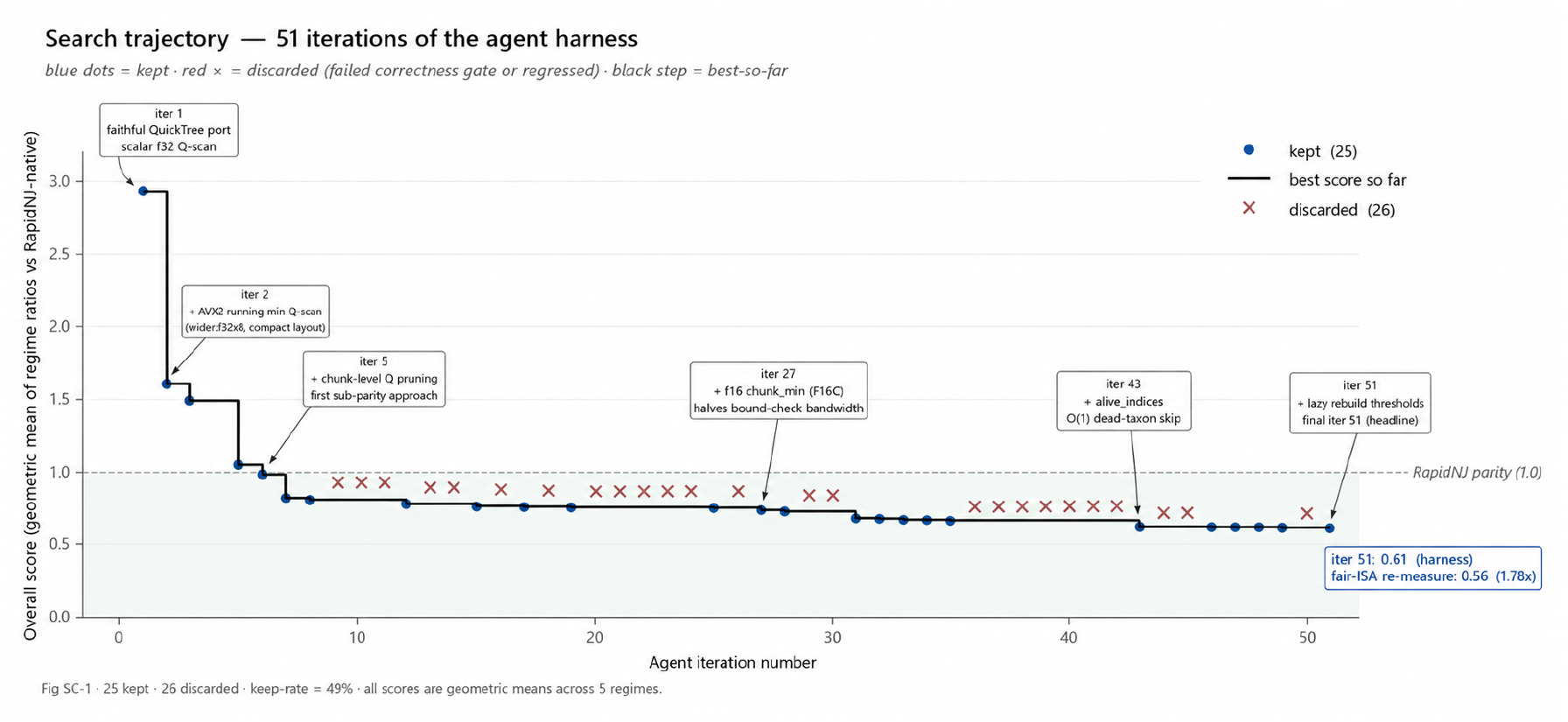
Search trajectory across the 51 iterations of the agent loop. Blue dots are kept iterations; red crosses are discarded for failing a correctness gate or regressing the score. The black step is best-so-far. Labelled milestones correspond to the five ingredients of Section 3.3.

## 5 Discussion

The result provides one empirical data point on the broader question of where current-generation coding agents are able to contribute, and under what oversight conditions, on tasks with established native-code baselines. Several properties of the target made it tractable. Exact NJ has a short kernel (a single *O*(*n*^3^) loop with auxiliary structures), a precise correctness oracle (RF distance against a reference tree), and cheap repeated measurement (seconds to minutes per matrix). These properties allowed each iteration to be evaluated within minutes rather than days, and allowed rejected proposals to be discarded without ambiguity.

We conjecture that the same combination of properties, short kernel, precise correctness oracle, and cheap measurement, identifies a class of bioinformatics targets where similar harness-gated approaches are likely to yield comparable constant-factor improvements. Candidate targets include maximum-likelihood phylogenetic inference, multiple sequence alignment, and several hot kernels in metagenomic analysis. The applicability of the approach to domains lacking a precise correctness oracle, or with expensive measurement, is open.

### 5.1 Boundaries of the result

This is a one-machine result on one benchmark corpus and one instruction-set family. Generalisation to AVX-512 hardware, to other CPU vendors, and to other compiler toolchains is not measured. The correctness gate at *n >* 2000 uses a branch-length tolerance check rather than RF, because QuickTree does not scale to those matrix sizes; this is a weaker form of correctness evidence than the RF-zero gate used at smaller *n*. The running agent was Opus 4.7 throughout; this paper does not run a controlled comparison against other models, against a non-agent baseline, or against the same model without the harness.

The harness is not released as part of this artefact, as it embeds task-specific prompt scaffolding and corpus paths that are not portable as-is. A reusable harness is a separate piece of work.

### 5.2 Iteration budget

The loop terminated at iteration 51 not because the running agent had exhausted productive proposals, the score trajectory was still descending and a labelled set of candidate proposals remained unevaluated, but because the rate limits on the personal Claude and Codex subscriptions used to run this project were reached. The optimisation may therefore be possible to push further under a less constrained compute budget. The same harness shape, run with dedicated compute over a longer time horizon, may yield additional improvement on this target; the present study does not test that conjecture. If it holds, it has practical implications for research groups whose hot computational kernels are stable, well-instrumented, and amenable to oracle-based correctness gates.

## 6 Conclusion

We presented SwiftNJ, a Rust implementation of exact neighbour joining produced by an LLM coding agent operating under a correctness-gated optimisation harness. SwiftNJ achieves a geometric-mean runtime ratio of 0.565 against a locally-rebuilt RapidNJ-native baseline on a 59-matrix benchmark and a 1.43*×* geometric-mean speedup on five large real-derived matrices outside the optimisation loop, with RF distance zero against the QuickTree reference on 400 shuffled inputs from 16 small matrices (*n ≤* 2000). The SwiftNJ source code is released under the MIT licence; benchmark inputs and generation recipes are released or cited with per-regime provenance and source terms at https://github.com/joe-b-christensen/swiftnj.

## Data Availability

The SwiftNJ source and all project-authored scripts and result data are released under the MIT licence. The synthetic regime (40 matrices) is regenerable from the shipped generator and seed manifest. The 14 Course matrices are Pfam/InterPro-derived protein-family distance matrices. The Pfam regime (5 matrices) is included and is also reproducible from public InterPro/Pfam (CC0) sources. The CRW 16S/23S matrices are cited but not shipped (size); the viral H1N1pdm/H5N1 matrices are reported as a transfer check only and are not shipped, because record-level INSDC provenance was not retained. Per-regime provenance, licences, and exact reproduction status are documented in `DATA_PROVENANCE.md`.

## Acknowledgements

I would like to thank Christian Nørgaard Storm Pedersen, Centre Director at the Bioinformatics Research Centre (BiRC), Aarhus University; this work began as a course assignment in his course, and the initial implementation built on ideas introduced there. The SwiftNJ Rust source was produced by an Opus 4.7 (xhigh) instance acting as the running agent inside the harness. The harness, the testing infrastructure, and the corpus curation were produced by me with assistance from two LLM collaborators (Opus 4.7 (xhigh) in Claude Code and GPT-5.4 (xhigh) in Codex).

